# *Foxi1* inactivation rescues loss of principal cell fate selection in *Hes1*-deficient kidneys but does not ensure maintenance of principal cell gene expression

**DOI:** 10.1101/826610

**Authors:** Malini Mukherjee, Jennifer DeRiso, Madhusudhana Janga, Eric Fogarty, Kameswaran Surendran

**Affiliations:** Sanford Research, 2301 East 60^th^ Street North, Sioux Falls, SD 57104, USA; Division of Basic Biomedical Sciences, Sanford School of Medicine, University of South Dakota, Vermillion, SD 57069; Department of Pediatrics, Sanford School of Medicine, University of South Dakota, Sioux Falls, SD 57104, USA

## Abstract

The distal nephron and collecting duct segments of the mammalian kidney consist of intercalated cell types intermingled among principal cell types. Notch signaling ensures that a sufficient number of cells select a principal instead of an intercalated cell fate. However, the precise mechanisms by which Notch signaling patterns the distal nephron and collecting duct cell fates is unknown. Here we observed that *Hes1*, a direct target of Notch signaling pathway, is required within the mouse developing collecting ducts for repression of *Foxi1* expression, an essential intercalated cell specific transcription factor. Interestingly, inactivation of *Foxi1* in *Hes1*-deficient collecting ducts rescues the deficiency in principal cell fate selection, overall urine concentrating deficiency, and reduces the occurrence of hydronephrosis. However, *Foxi1* inactivation does not rescue the reduction in expression of all principal cell genes in the Hes1-deficient kidney collecting duct cells that select the principal cell fate. Additionally, suppression of Notch/Hes1 signaling in mature principal cells reduces principal cell gene expression without activating *Foxi1*. We conclude that Hes1 is a Notch signaling target that is essential for normal patterning of the collecting ducts with intermingled cell types by repressing *Foxi1*, and for maintenance of principal cell gene expression independent of repressing *Foxi1*.

## Introduction

The collecting ducts of the mammalian kidney are composed of principal and intercalated cell types that ensure normal water, electrolyte and pH homeostasis. The ureteric bud grows and repeatedly branches to form the collecting duct system that connects all the nephrons to the ureter [1]. The majority of the ureteric bud cells mature into principal cells that express Aquaporin-2 (Aqp2) apically and express Aquaporin-3 (Aqp3), Aquaporin-4 (Aqp4) and Arginine vasopressin receptor 2 (Avpr2) on the basolateral membrane and are responsive to arginine vasopressin peptide [2]. Intermingled among the principal cells are the intercalated cells which express carbonic anhydrase II (CAII, encoded by *Car2*), specific subunits of the vacuolar-type H^+^-ATPases (v-ATPases), such as b1 (encoded by *Atp6v1b1*) and a4 (encoded by *Atp6v0a4*), and specific anion exchangers (AE) and play a critical role in systemic acid-base homeostasis [3-5]. The intercalated cells are classified into type A, type B and non-A non-B, based on the differential localization of v-ATPase proton pump and the type (AE1 versus AE4 or pendrin) and localization of AEs (apical versus basolateral) [6, 7]. The intercalated cells belong to a type of epithelia that are termed ionocytes that specialize in proton or bicarbonate secretion, and are present in a few different mammalian organ systems, and in frog and zebrafish skins [8-10]. Apart from the collecting ducts, the terminal part of distal convoluted tubules (DCT2) and connecting segment (CNT) also are composed of intercalated cell types [11].

Foxi1 and Notch signaling play a conserved role in regulating the differentiation of the proton or bicarbonate secreting epithelial cells, where the intercalated cell differentiation is promoted by *Foxi1*-orthologs and the fate selection is negatively regulated by Notch signaling in frogs, zebrafish and mice [10, 12-14]. Studies in mice have determined that inactivation of *Foxi1* results in distal renal tubular acidosis due to the absence of intercalated cell differentiation [13], and inactivation of different Notch signaling pathway components in the developing ureteric ducts results in nephrogenic diabetes insipidus-like phenotype due to increased number of ureteric duct cells differentiating into intercalated cells instead of principal cells [14, 15]. Notch signaling is considered to occur between adjacent mammalian cells, with one or more of the four mammalian Notch receptors being activated by ligands belonging to the Delta-like (Dll) and Jagged (Jag) family of type I transmembrane proteins [16]. The Notch receptors are serially cleaved upon ligand binding to release the Notch intracellular domain (NICD) from the membrane [17, 18]. NICD translocates to the nucleus, interacts with a DNA-binding factor RBPJ and recruits mastermind-like to activate target genes, such as the transcriptional repressors Hairy/Enhancer of Split (HES) family members Hes1 and Hes5 [19].

Here we wished to further understand the mechanisms by which Notch signaling regulates the collecting duct cell fate selection and differentiation. In mice with Notch-signaling-deficient ureteric bud, the number of Foxi1^+^ (intercalated) cells is increased, while the number of principal cells is reduced [14, 15, 20]. Foxi1 is a transcription factor specifically expressed in the intercalated cells of the kidneys, is necessary for intercalated cell differentiation [13], and it activates the expression of intercalated cell specific genes such as *Atp6v1b1, Atp6v0a4, Slc26a4*, and *Slc4a9* [21-23]. Based on these observations, along with knowledge of how Notch signaling mediates cell fate selection in different developmental settings in which neighboring cells take on different cell fates [24-26], it is hypothesized that Notch receptor activation mediates a lateral inhibitory signal to repress an essential intercalated fate promoting transcription factor, such as *Foxi1* expression, to allow for the principal cell program to be turned on (Fig.1). However, the precise mechanism by which Notch signaling represses the intercalated cell fate remains to be determined. We have previously observed that ectopic expression of activated Notch1 in the developing collecting ducts activates principal cell-specific genes such as *Elf5* prior to repressing *Foxi1* [20]. This opens the possibility that Notch signaling represses an up stream activator of *Foxi1* to prevent intercalated cell fate selection which in turn represses *Elf5* expression, or that Notch signaling can directly activate expression of some principal cell specific genes independent of repressing the intercalated cell fate selection. In the current study we examine whether Notch signaling promotes any aspect of the principal cell program independent of repressing *Foxi1* expression.

**Figure 1.**
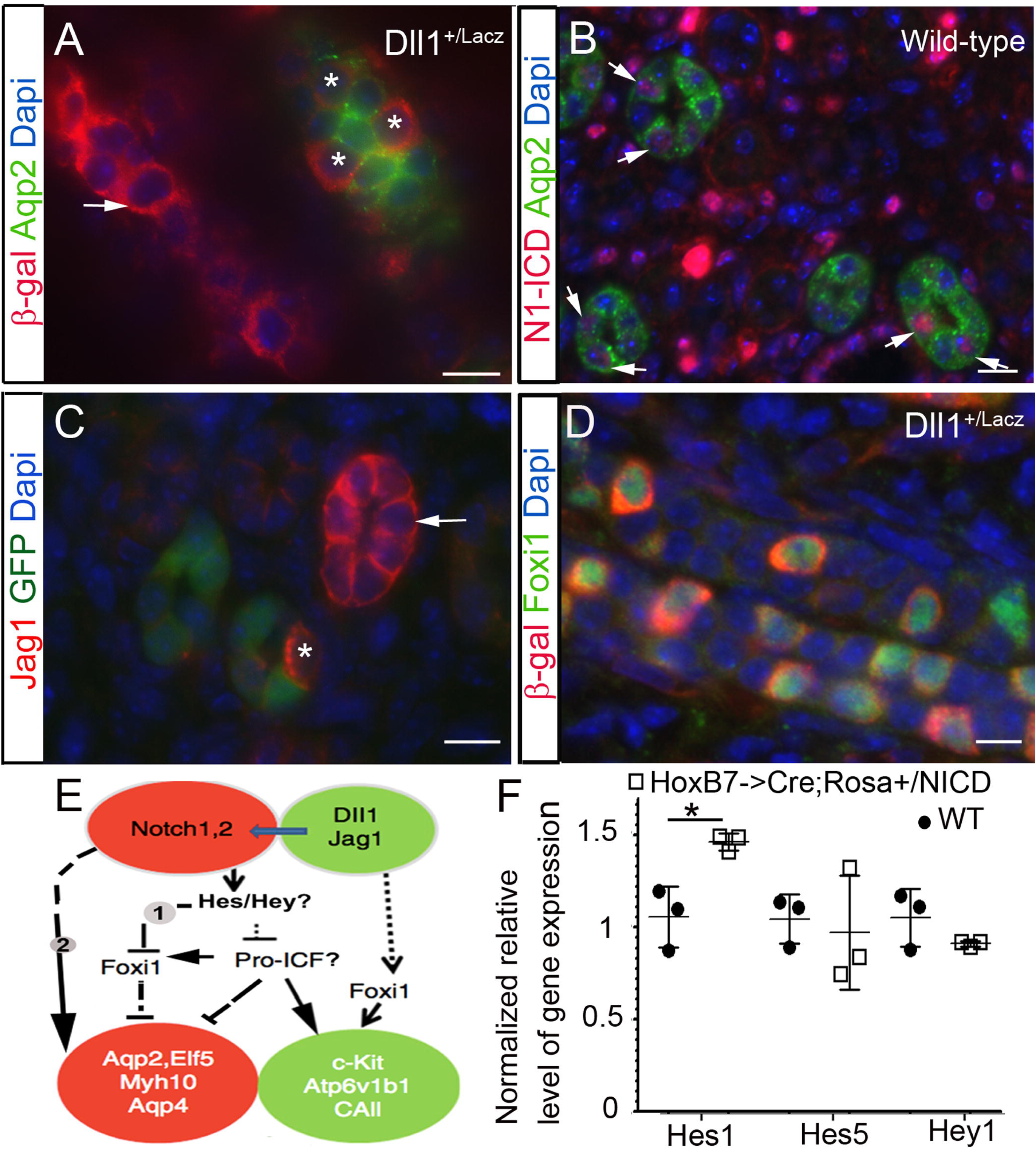
Notch signaling within the collecting duct epithelium patterns the duct cell fates by unknown mechanisms. **A.** The Notch ligand Delta-like 1 (Dll1) is expressed in cells (asterisks) adjacent to Aqp2^+^ principal cells as revealed by staining for β-galactosidase (□-gal) in *Dll1*^*+/LacZ*^ [29] in E16.5 kidneys. **B**. Activated Notch1 (N1-ICD) is detected in Aqp2^+^ cells (arrows) of E17.5 mouse kidneys. **C**. Another Notch ligand, Jagged1 (Jag1), is expressed in a cell (asterisk) adjacent to *Elf5*-expressing principal cells as revealed by GFP expression (a surrogate for *Elf5*) in the kidneys of E18.5 *Elf5->GFP* transgenic mice. **D**. β-gal as a surrogate for *Dll1* is strongly expressed in the Foxi1-expressing cells in P0 kidneys of *Dll1*^*+/LacZ*^ mice. **E**. Possible mechanisms of how CD cell fate patterning may be mediated by Notch signaling. Notch receptors are activated in cells that will become PCs, while *Dll1* and *Jag1* are strongly expressed in cells that will become intercalated. The UB-derived cells turn on Foxi1 and become ICs. Activated Notch receptors inhibit the expression of Foxi1 in UB cells by an unknown mechanism-1 to allow for principal cell gene expression. Additionally, Notch signaling may independent of repressing *Foxi1*, activate expression of principal cell genes, such as *Elf5* by mechanism-2. **F**. Ectopic expression of activated N1-ICD in the developing kidney collecting ducts increases the expression of Hes1 but not Hes5 or Hey1 in E14.5 *HoxB7->Cre;Rosa*^*+/NICD*^ mouse kidneys when compared with wild-type littermate kidneys (n=3 per genotype), asterisk denotes p<0.05, two-tailed unpaired t-test. Scale bars are 10μm.

## Methods

### Mice

All experiments involving mice were approved by the Sanford Research IACUC. Details of mouse lines used are listed in Supplementary Table1. Mice used in this study were maintained on mixed backgrounds and genotyped following a universal PCR genotyping protocol [27]. Age and sex matched control littermates were housed together and we examined mice of both sexes. For timed mating mice were observed daily for the presence of vaginal plugs and noon on the day a plug was observed was considered E0.5. Primer sequences are available upon request.

### Histology and immunohistochemistry

Kidneys were fixed in 4% PFA or Bouin’s fixative overnight at 4°C, washed in 70% ethanol, paraffin embedded and sectioned at 12mm thickness. Prior to immunostaining, the sections were de-paraffinized in xylene, rehydrated and, boiled for 20 min in Trilogy (Cell Marque) for antigen unmasking. Sections were blocked in PBS containing 1% bovine serum albumin (BSA), 0.2% powdered skim milk, and 0.3% Triton X-100 for at least 15 min at RT prior to incubation with primary antibodies overnight. For direct visualization of GFP or tdtomato the mouse kidneys were fixed in 4% PFA for 1 hour, rinsed in PBS, and incubated in 15% sucrose for 24 hours and then another 24 hours in 30% sucrose prior to embedding in OCT and sectioning at 12mm thickness. Primary antibody details are listed in Supplementary Table 2.

### RT-qPCR: Taq-man and SYBR green

RNA from whole kidneys were extracted using commercial RNA mini or midi kits (Qiagen). RNA was reverse transcribed using random hexamers or oligo dT with reverse transcription kit (Promega). Quantitative PCR was performed using Power SYBR Green (Life Technologies), gene specific primers with the forward and reverse primers designed from different exons, and an ABI 7500 instrument (Applied Biosystems). Standard curves were generated analyzing serially diluted cDNA reverse transcribed from mouse kidneys to determine the efficiency of each primer pair. Each sample was measured either in duplicate or triplicate and relative gene expression levels were normalized to that of *GAPDH* or *beta-2 microglobulin*. For detecting changes in genes that have low abundance, 5µg RNA was reverse transcribed using gene specific primer, and the resultant cDNA was diluted minimally to perform qPCR assays.

### Cell culture

We utilized the mature principal cell line mpkCCDc14 from Dr. Vandewalle [28]. For generating stable clones, cells were transfected with expression plasmid of dnMamL and selected for neomycin resistance.

### Statistics

Initial studies of three mouse kidneys with ectopic expression of NICD and three wild-type littermate kidneys revealed these numbers were sufficient to detect significant differences between the two groups by two-tailed t-test, with α=0.05 and Power=0.8, resulting in an effect size of 2.4 or higher. We therefore continued the studies with n=3 or more per group with the exact numbers depending on the size of and genotype within the litters. For urine osmolality the sample size also depended on whether or not a sufficient volume of urine was collected. In the graphs, the height of each bar represents the mean and the error bars represent one standard deviation. In the scatter plots the large horizontal line through a group of data points represents the mean and the smaller horizontal lines flanking the mean in each group represent one standard deviation. Excel was used to perform two-tailed unpaired t-tests to compare two groups of mice or cells, after verification that samples had a normal distribution and testing for equal variance between groups using the F-test. The resulting p values are stated in the text and figure legends. For urine osmolality in combined Foxi1 and Hes1 mutants, a two-way ANOVA was performed followed by multiple pair-wise comparison of the different genetic groups using the Tuckey HSD test.

## Results

### The Notch-ligands Dll1 and Jag1 are expressed in intercalated cells adjacent to principal cells and ectopic expression of activated-Notch1 increases the expression of the transcriptional repressor Hes1

The hypothesis that Notch signaling represses *Foxi1* would predict that the Notch ligands are expressed in the intercalated cells, while the Notch receptors are activated in principal cells. The expression of the ligand *Dll1* reported by β-galactosidase in *Dll1*^*+/LacZ*^ mice [29] at E16.5 shows that the developing collecting duct cells that have selected the principal cell fate as determined by Aqp2 expression are adjacent to duct cells with the highest *Dll1* expression (asterisks in Fig.1A), while activated Notch1 (N1-ICD) is observed in E17.5 Aqp2^+^ duct cells (arrows in Fig.1B), and Jagged-1 (Jag1), another Notch ligand, is expressed in duct cells adjacent to cells expressing GFP in E18.5 *Elf5->GFP* transgenic mouse kidneys (Fig.1C). We have previously determined that Elf5 is dependent on Notch signaling and is an early principal cell lineage specific transcription factor that can activate *Aqp2* and *Avpr2* expression [20]. At birth all the collecting duct cells with high level of *Dll1* expression also express Foxi1, an intercalated cell marker (Fig.1D). The expression patterns of Notch-ligands, N1-ICD and Notch-regulated gene, *Elf5*, are suggestive that developing collecting duct cells expressing the Notch-ligands activate Notch signaling in the adjacent cells to promote principal cell differentiation (Fig.1E).

Based on the *Dll1*, Jag1 and N1-ICD expression patterns, Notch signaling may simply inhibit *Foxi1* and other still to be identified intercalated cell factors (pro-ICFs) to prevent intercalated cell fate selection (Fig.1E, pathway-1). In this model, Foxi1 and/or other ICFs expressed in the ligand-expressing cells inhibit *Elf5* and other principal cell-specific transcription factors to ensure that the principal cell program is not selected in intercalated cells. Alternatively, in addition to inhibiting *Foxi1* and additional ICFs, Notch-signaling may also more directly turn on the principal cell program by a mechanism not dependent on inhibiting *Foxi1* (Fig.1E, pathway-2). How Notch signaling mediates repression of *Foxi1* and activation of principal cell specific genes such as *Elf5* remains unknown. Since Hairy/Enhancer of Split (Hes) genes code for transcriptional repressors that function as downstream targets of Notch signaling we examined the expression of Hes family of genes following ectopic expression of N1-ICD in the developing collecting duct which we previously observed promoted increased principal cell gene expression and increased expression of *Hes1*, a known Notch-target gene in the E13.5 collecting ducts [20]. Here we confirmed that ectopic expression of N1-ICD in the developing collecting duct increases *Hes1* expression at E14.5, and did not alter *Hes5* or *Hey1* expression levels (Fig.1F) which are known be expressed in the developing kidneys [30, 31]. Considering that *Hes1* codes for a transcriptional repressor that is activated by N1-ICD and is expressed in the developing collecting ducts [31], it may mediate the repression of *Foxi1* during kidney collecting duct development. However, a previous study reported that E16.5 kidneys from *Hes1*^−/−^; *Hes5*^*+/-*^ mice appeared normal [31], and hence it is necessary to determine whether Hes1 regulates kidney collecting duct cell type patterning during development.

### Hes1 is required for ensuring the development of normal principal to intercalated cell ratio and repression of *Foxi1* in the developing kidney

To test if Hes1 functions down-stream of Notch signaling during collecting duct development to mediate the repression of *Foxi1* we inactivated *Hes1* floxed alleles (Fig. 2A) in *Hes1*^*f/f*^ mice [32] using *Cdh16->Cre* transgene which drives Cre expression in the distal tubules and collecting ducts [33]. We verified that the *Hes1* floxed allele is recombined in the kidneys but not in tail specimens of *Cdh16->Cre* inheriting mice (Fig. 2B). The post-natal day 0 (P0) kidneys of *Cdh16->Cre;Hes1*^*f/f*^ mice have an obvious increase in CAII-expressing cells and a decrease in Aqp2-expressing cells (Fig. 2G-J) when compared with that of wild-type littermates (Fig. 2C-F). This change in principal (Aqp2^+^) to intercalated (CAII^+^) cell ratio in *Cdh16->Cre;Hes1*^*f/f*^ mouse kidneys compared with wild-type littermates is most evident in the cortex (Fig.2C and G) and outer medulla (Fig.2D and H) and subtle in the inner medulla (Fig.2E and I) and papilla (Fig.2F and J). Additionally, there are many duct cells that are negative for both Aqp2 and CAII in the *Cdh16->Cre;Hes1*^*f/f*^ mouse kidneys, especially in the papillary regions (arrows in Fig.2J). To further validate the change in cell ratio we quantified the expression levels of principal cell specific genes by RT-qPCR. We observed reduced mRNA levels of principal cell specific genes *Elf5, Avpr2, Aqp2* and *Aqp4*, along with reduced *Hes1* mRNA levels in *Cdh16->Cre;Hes1*^*f/f*^ mouse kidneys compared with that of wild-type littermates (Fig. 2K). Consistent with a reduction in principal cell numbers the *Cdh16->Cre; Hes1*^*f/f*^ mice (n=17) have a significantly reduced urine concentrating capacity as on average the urine osmolality is reduced by 50% when compared with wild-type littermates (n=8) at 1 month of age (Fig. 2L). Hence, Hes1 is critical for principal cell fate selection, and may additionally ensure normal expression of Aqp2 following cell fate selection. Apart from the reduction in principal cell specific gene expression, *Foxi1* expression is increased in *Cdh16->Cre; Hes1*^*f/f*^ mouse kidneys (Fig. 2k). These observations are consistent with Hes1 mediating Notch signaling dependent principal cell fate selection by repression of *Foxi1* expression.

**Figure 2.**
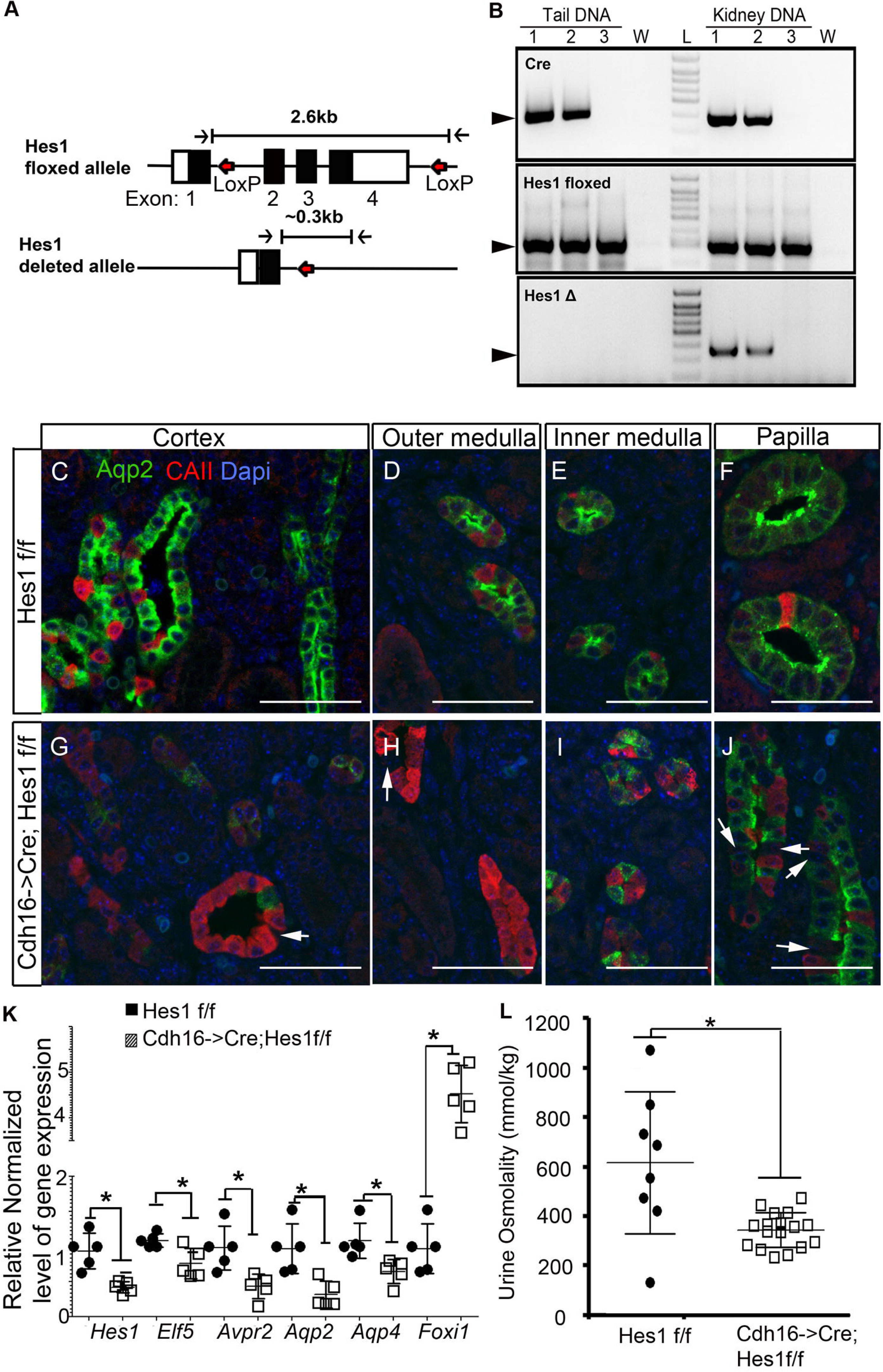
The Notch target Hes1 is required for ensuring normal patterning of collecting ducts with principal and intercalated cells. **A**. Schematic of Hes1 floxed allele with LoxP sites flanking exons 2 to 4 and Hes1 deleted allele missing exons 2 to 4 following cre mediated LoxP recombination. **B**. Genotyping using genomic DNA extract from tail and kidney confirm kidney specific recombination of LoxP sites in mice that inherited Cdh16->Cre (lanes 1 and 2) but not in mice without Cre (lane 3). W = water, L= DNA ladder. **C-J** Staining for Aqp2 and CAII reveals that inactivation of *Hes1* in the developing kidney (G-J) leads to an increase in intercalated cells and a decrease in principal cell numbers compared to control littermates (C-F). This change in principal (Aqp2^+^) to intercalated (CAII^+^) cell ratio is most evident in the cortex (C and G) and outer medulla (D and H) and subtle in the inner medulla (E and I) and papilla (F and J). Arrows point at cells negative for both Aqp2 and CAII. **K**. Consistent with a change in the ratio of principal to intercalated cell types there is a decrease in gene expression of principal cell-specific genes and an increase in intercalated gene *Foxi1* (* denotes p<0.05, two-tailed unpaired t-test; n=5 per genotype). **L**. Mice with Hes-1 deficient collecting ducts have significantly reduced urine concentrating capacity compared with wild-type littermates (* denotes p<0.05, two-tailed unpaired t-test; n=8 for wild type and n=17 for *Cdh16->Cre; Hes1*^*f/f*^ mice).

### Inactivation of *Foxi1* is sufficient to rescue principal cell fate selection deficiency in the absence of Hes1

To determine if Notch signaling via Hes1 simply inhibits *Foxi1* expression to prevent Foxi1-mediated repression of the principal cell program, we compared *Cdh16->Cre; Hes1*^*f/f*^ mice with and without *Foxi1* inactivation. If repression of *Foxi1* is the main mechanism by which Notch signaling promotes the principal cell fate selection, then we expect that inactivation of *Foxi1* should rescue the principal cell deficiency in *Cdh16->Cre; Hes1*^*f/f*^ mice. We analyzed P0 kidneys from *Cdh16->Cre; Hes1*^*f/f*^; *Foxi1*^*+/\**^ (Hes1-mutant, * denotes + or – allele), *Foxi1*^−/−^; *Hes1*^*\*/f*^ (Foxi1-mutant, * denotes + or f allele), *Cdh16->Cre; Hes1*^*f/f*^; *Foxi1*^−/−^ (Hes1&Foxi1-double mutant) and *Hes1*^*f/f*^; *Foxi1*^*+/\**^ (wild-type littermates, * denotes + or – allele). We stained for Aqp2, which is normally present in cytoplasmic vesicles and transported to apical membrane of principal cells in response to arginine vasopressin peptide (AVP), and for Aqp4, which localizes to the basolateral membranes of principal cells. These principal cell markers turn on in the stalk of the branching ureteric duct and are absent in the duct tips (Fig. 3A-D) [20]. Interspersed among the principal cells are Aqp2^−^ and Aqp4^−^ and likely intercalated cells in wild type kidney (arrowheads in Fig.3E, I & M). Apart from an increase in number of intercalated cells in Hes1-mutant mouse kidneys (Fig.2G-I, and arrowheads in Fig.3F, J & N), we observed several principal cells that are Aqp4^+^ but do not express Aqp2 (arrows in Fig. 3F, J and N). There are some collecting duct segments (asterisks in the lumen in Fig.3F &J) in the Hes1-mutant mouse kidneys that completely lack Aqp2 expression but retain reduced levels of Aqp4 expression (Fig. 3F&J). These observations are suggestive that apart from Hes1 being critical for principal cell fate selection, Hes1 is also required for ensuring normal levels of Aqp2 and Aqp4 expression following principal cell fate selection. In the Foxi1-mutant P0 kidneys, apart from the collecting duct cells near the duct tips and proximal duct stalk regions which normally have not turned on Aqp2 or Aqp4 expression, the Aqp2^−^; Aqp4^−^ intercalated cells are absent in the cortex and medulla (Fig.3C,G&K). Additionally, we were not able to detect Atp6v1b1 or AE1 protein expression within the epithelial structures of Foxi1 mutant mouse kidneys by immunohistochemistry (Fig.S1). These observations are consistent with Foxi1 being critical for intercalated cell development [13]. Additionally, prior characterization of Foxi1-mutant kidneys suggest that Foxi1-deficient collecting duct cells arrest at an intermediary stage since they were observed to express both Aqp2, a principal cell marker, and CAII, an intercalated marker [13]. Our analysis revealed that the P0 Foxi1-mutant kidneys do not express detectable levels of CAII in the collecting ducts (Fig.S1). However, at 7 weeks the Foxi1-mutant kidneys do show extensive co-expression of CAII in Aqp2 expressing cells (Fig.S1). Additionally, if the Foxi1-deficient collecting ducts arrest at an intermediary state we expected to observe Aqp2^+^; Aqp4^−^ cells in the Foxi1-mutant kidneys, assuming that Aqp4 is a mature principal cell marker, instead we observed that most collecting duct cells are Aqp2^+^; Aqp4^+^. We did observe Aqp2^**-**^; Aqp4^**-**^ cells in the papillary regions of Foxi1-mutant kidneys (arrowheads in Fig.3O) similar to that in wild-type kidneys (arrowheads in Fig.3M). These observations are suggestive that in the absence of Foxi1 most duct cells select the principal cell fate, and CAII turns on in the principal-like cells of adult Foxi1 null mouse kidneys.

**Figure 3.**
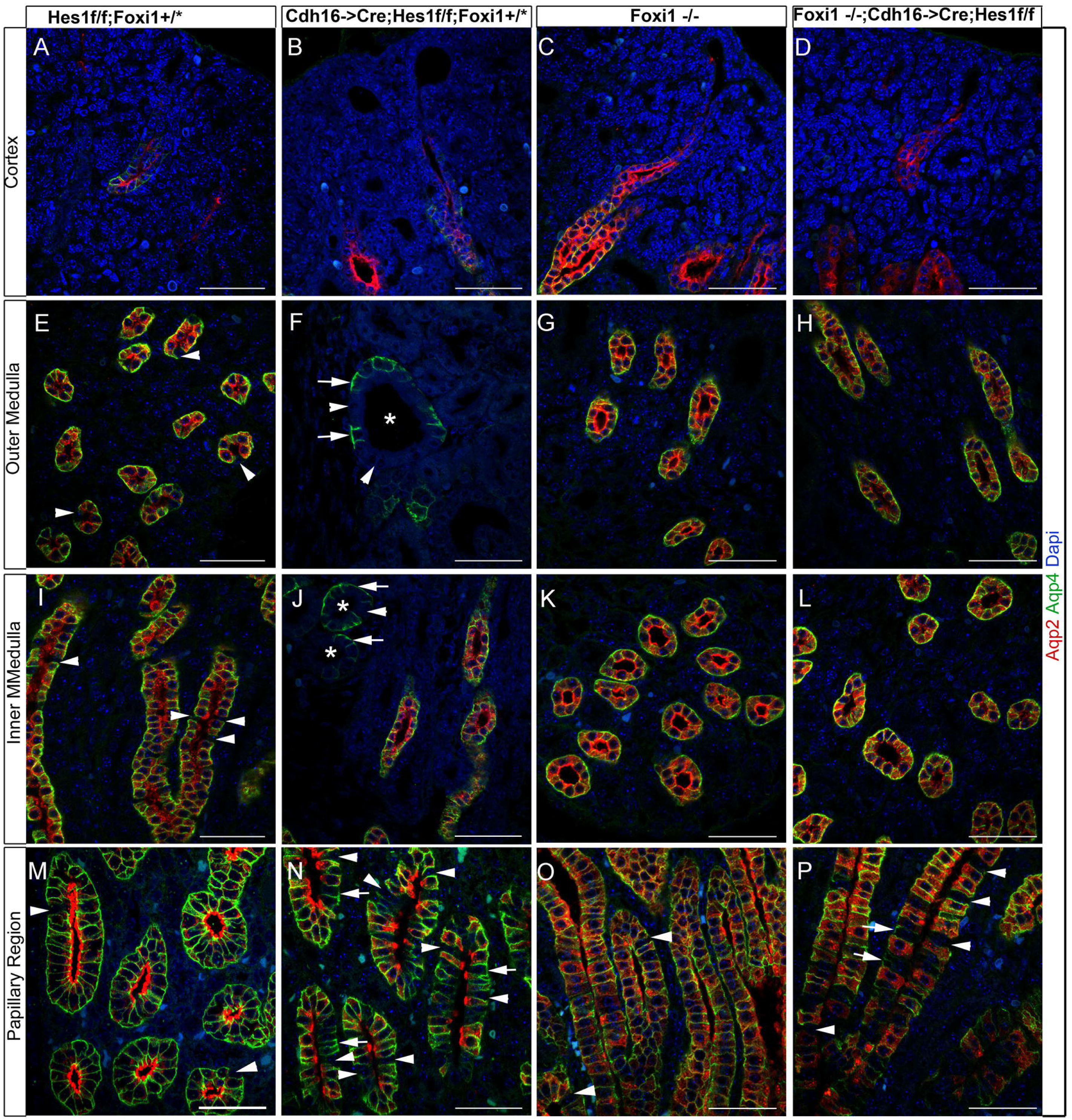
Staining for Aqp2 and Aqp4 reveals that inactivation of *Foxi1* in *Cdh16->Cre; Hes1*^*f/f*^ mice recuses the principal cell fate selection deficiency. A, E, I &M. Wild type neonatal mouse kidneys show expected ratios of principal to intercalated cells (arrowheads in E, I & D) in collecting duct segments. **B, F, J &N.** Inactivation of *Hes1* results in dilated collecting ducts (asterisk in F&J) with no Aqp2 expression in some principal cells (arrows in F, J & N) along with a reduction in number of principal cells. **C, G, K&O.** Foxi1-deficient mice develop collecting ducts with increased number of cells expressing markers of principal cells. **D, H, L&P** *Foxi1* inactivation rescues Hes1-deficiency-dependent defects in principal cell fate selection in all regions expect the papillary regions where many intercalated (Aqp2^−^; Aqp4^−^) cells remain (arrowheads in P). Additionally, the reduction in Aqp2 and Aqp4 expression levels in *Hes1*-null principal cells (arrows in J and N) is not restored by *Foxi1* inactivation in some principal cells within the papillary regions (arrows in P). Scale bars are 50μm.

Interestingly, inactivation of *Foxi1* in the Hes1-deficient collecting ducts rescues the principal cell fate selection deficiency observed in Hes1-mutant mice in most regions of the kidney as we observe increased number of Aqp2^+^; Aqp4^+^ cells in the Hes1&Foxi1-double mutant mouse kidneys (Fig. 3). The papillary regions of Hes1&Foxi1-double mutants (arrowheads in Fig. 3P) still have many more Aqp2^**-**^; Aqp4^**-**^ cells when compared to wild-type papillary regions (arrowheads in Fig. 3M) and to Foxi1-mutant papillary regions (arrowheads in Fig. 3O). Inactivation of *Foxi1* also does not rescue the reduced levels of Aqp2 and Aqp4 expression in some principal cells within the papillary region of Hes1&Foxi1-double mutants (arrows in Fig. 3P). Overall, these observations are suggestive that while Hes1 mediated repression of *Foxi1* is sufficient for principal cell fate selection, there may be additional functions for Hes1 that are independent of Foxi1 in ensuring normal expression levels of Aqp2 and Aqp4 within principal cells after cell fate selection.

### *Foxi1* inactivation rescues the Hes1-deficiency dependent urine concentrating deficit, hydronephrosis and the reduced expression of some but not all principal cell genes

Considering that *Foxi1* inactivation rescues the principal cell fate selection deficiency observed in kidneys of *Cdh16->Cre; Hes1*^*f/f*^ mice at birth, we next examined whether this was sustained over time and if it was sufficient to allow for normal urine concentrating capacity. Staining for principal cell markers: Aqp2, Aqp3 and Aqp4 along with the intercalated maker c-Kit in 7 week old kidneys revealed that *Cdh16->Cre; Hes1*^*f/f*^ mice continued to have reduction in principal cells and an increase in intercalated cells (Fig. S2). Interestingly, inactivation of *Foxi1* in the *Cdh16->Cre; Hes1*^*f/f*^ mice continues to rescue the deficiency in principal cell numbers (Fig.S2 & Fig.4E). Analysis of urine osmolality revealed a significant deficiency in urine concentrating capacity in *Cdh16->Cre;Hes1*^*f/f*^ mice in two separate cohorts of mice analyzed at 4 and 7 weeks of age compared with wild type littermate controls (Fig. 4A). Interestingly, inactivation of *Foxi1* in *Cdh16->Cre;Hes1*^*f/f*^ mice significantly restores the urine osmolality (Fig.4A). Two way ANOVA showed that the interaction between age and genetic modification is not significant. As a result, this interaction effect was left out in modelling the data. Both the genetics and age, as independent factors, have significant effects on urine osmolality. Overall, the urine osmolality was higher in collections at 7 weeks of age compared to 4 weeks of age. However, the *Cdh16->Cre;Hes1*^*f/f*^ mice had a lower than normal urine osmolality at 4 weeks and did not show a further increase at 7 weeks. Multiple pair-wise comparisons of the urine osmolality of the different genetic groups was conducted using the Tuckey-HSD test and revealed a significant difference between Wild type (WT) and *Cdh16->Cre;Hes1*^*f/f*^ (Hes1-mutant), as well as between Hes1-mutant and *Cdh16->Cre;Hes1*^*f/f*^ *;Foxi1*^−/−^ (Hes1 & Foxi1 mutant). Consistent with previous reports of mice with defective principal cell function developing hydronephrosis, we observed hydronephrosis resulting in renal medullary cavities in at least one kidney in 75% of *Cdh16->Cre;Hes1*^*f/f*^ mice at 1 to 3 months of age (Fig. 4B &C). *Foxi1* inactivation in Hes1-mutant mice reduced the occurrence of hydronephrosis to 25% (Fig.4B&D).

**Figure 4.**
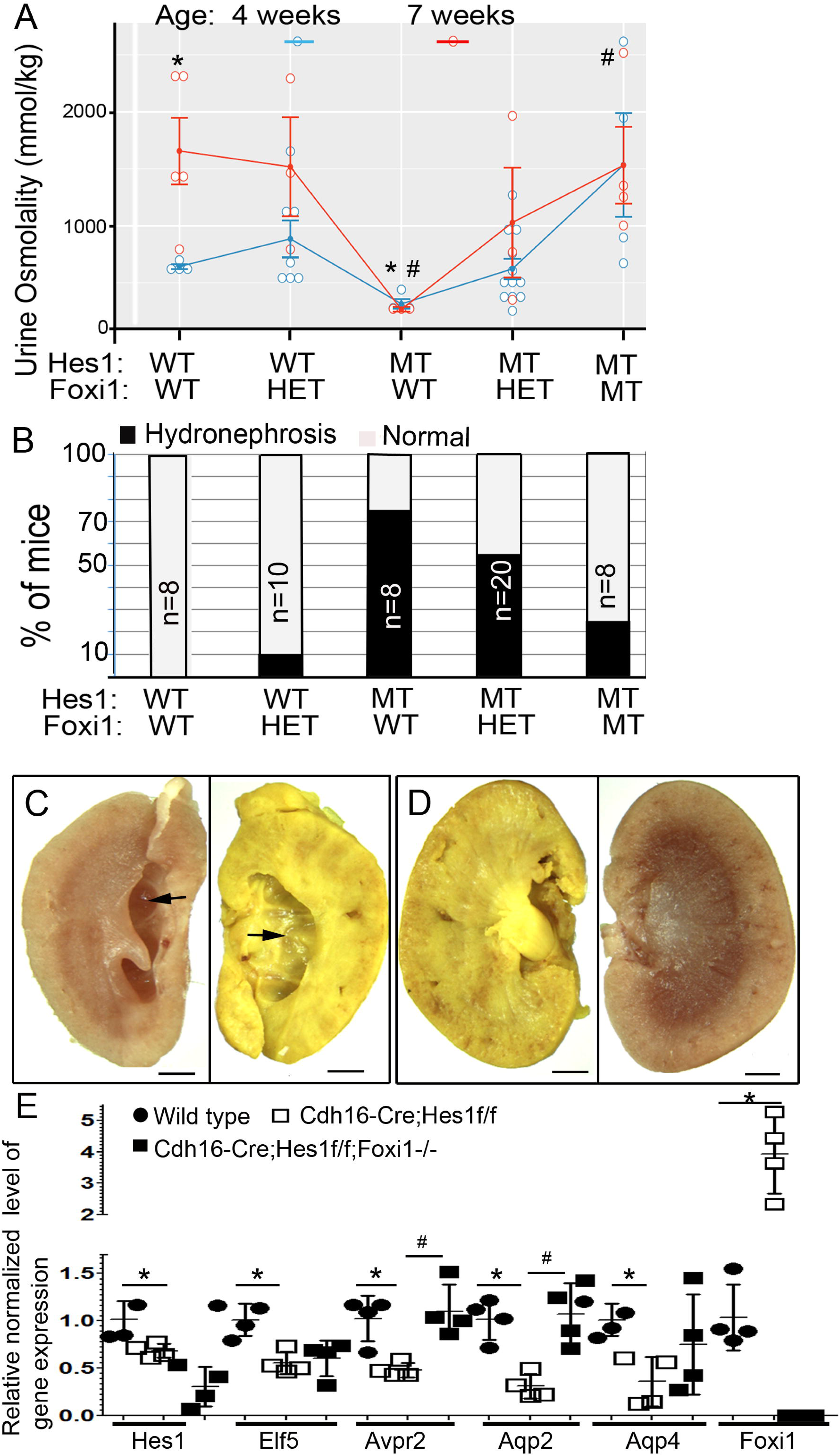
*Foxi1* inactivation rescues the Hes1-deficiency dependent urine concentrating deficit, hydronephrosis and the reduced expression of some but not all principal cell genes. **A.** Graph comparing the urine osmolality of mice with *Hes1* and/or *Foxi1* inactivation at 4 and 7 weeks of age. WT = wild type, HET= heterozygous, and MT= mutant. ANOVA followed by multiple pair-wise comparisons of the different genetic groups using the Tuckey-HSD test revealed a significant difference between Wild type (WT) and *Cdh16->Cre;Hes1*^*f/f*^ (Hes1-mutant) urine osmolality, denoted by asterisks. Similarly, there is a significant difference between Hes1 mutant and *Cdh16->Cre;Hes1*^*f/f*^ *;Foxi1*^−/−^ (Hes1 & Foxi1-mutant) urine osmolality, denoted by #. **B.** Graph depicting the percentage of mice within each genetic group that develop hydronephrosis in at least one kidney. **C & D**. Images of bisected kidneys from *Cdh16->Cre;Hes1*^*f/f*^ mouse (C) reveals medullary cavity formed due to hydronephrosis (arrows), which is rescued in *Cdh16->Cre;Hes1*^*f/f*^ *;Foxi1*^−/−^ mice (D). **E.** Relative normalized gene expression levels determined by RT-qPCR of whole kidney RNA reveals that Hes1 and genes expressed in principal cells are significantly down-regulated in Hes1-mutant kidneys compared with wild type kidneys (asterisks denote p<0.05, n=4 per group, two-tailed unpaired t-test). *Foxi1* inactivation in mice with Hes1-deficient collecting ducts rescues reduced *Avpr2* and *Aqp2* expression levels in mice with Hes1-deficient collecting ducts (# denotes p<0.05, n=4 per group, two-tailed unpaired t-test).

To further determine whether *Foxi1* inactivation recuses Hes1-mutant defects we examined the expression levels of principal cell specific genes in wild-type, versus Hes1-mutant versus Foxi1 & Hes1-mutant kidneys of 4 week old mice by RT-qPCR. First we confirmed that *Hes1* levels are reduced in the *Cdh16->Cre;Hes1*^*f/f*^ and *Cdh16->Cre;Hes1*^*f/f*^; *Foxi1*^−/−^ kidneys compared with wild type littermates, and *Foxi1* levels are reduced in *Cdh16->Cre;Hes1*^*f/f*^; *Foxi1*^−/−^ kidneys (Fig. 4E). Then we confirmed that genes expressed in principal cells including *Elf5, Avpr2, Aqp2* and *Aqp4* are significantly down regulated in Hes1-mutants compared with wild type controls (asterisks in Fig.4E). Among the analyzed principal cell genes, only the expression levels of *Aqp2* and *Avpr2* returned to wild type levels with *Foxi1* inactivation in the Hes1-deficient kidneys. Both Aqp2 and Avpr2 expression levels are significantly increased in Foxi1&Hes1-mutants compared with Hes1-mutants (# in Fig.4E). While *Elf5* and *Aqp4* expression levels are reduced in Hes1-mutant kidneys the expression levels of these principal genes were not rescued to normal levels in *Cdh16->Cre;Hes1*^*f/f*^; *Foxi1*^−/−^ kidneys. Consistent with *Foxi1* inactivation not being able to rescue the expression level of all principal cell specific genes we observed a reduction in basolateral Aqp4 protein expression in some collecting duct segments of *Cdh16->Cre;Hes1*^*f/f*^; *Foxi1*^−/−^ mouse kidneys (Fig.S2F). In summary, most of the kidney defects observed in *Cdh16->Cre;Hes1*^*f/f*^ mice such as reduced principal cell numbers, reduced urine osmolality and hydronephrosis are all rescued with the additional inactivation of *Foxi1*. However, not all principal cell gene expression is restored to normal levels with the inactivation of *Foxi1* in Hes1-mutant kidneys.

### Suppression of Notch signaling in mature principal cells down regulates expression of principal cell specific genes without activating *Foxi1* expression

The reduced expression of Aqp2 and Aqp4 protein in some principal cells in both *Cdh16->Cre;Hes1*^*f/f*^ and *Cdh16->Cre;Hes1*^*f/f*^ *;Foxi1*^−/−^ kidneys is suggestive that Hes1 is required for maintenance of normal level of principal cell gene expression following principal cell fate selection. Additionally, the inability of *Foxi1* inactivation to rescue the expression levels of *Aqp4* and *Elf5* mRNA in *Cdh16->Cre;Hes1*^*f/f*^ *;Foxi1*^−/−^ kidneys implies that Hes1 functions in a Foxi1 independent manner to maintain principal cell gene expression. To being determining the mechanism by which Notch signaling regulates principal cell specific gene expression we established three established three independent stable clones of the mature principal kidney cortical collecting duct (mpkCCDc14) cells [28] expressing dominant-negative mastermind-like-1 (dnMaml). Consistent with dnMaml functioning as a repressor of Notch signaling mediated transcription we observed reduced expression of *Hes1* in the principal cells expressing dnMaml (Fig. 5A). Along with the down regulation of *Hes1* mRNA levels we observed reduced expression of principal cell genes *Aqp2, Aqp4, Avpr2* and *Elf5* (Fig. 5A). Interestingly, the down regulation of principal cell genes occurs without increased expression of *Foxi1* in the mpkCCDc14 cells expressing dnMaml (Fig.5B). These observations indicate that de-repression of *Foxi1* is not necessary for down-regulation of principal cell specific genes in mature principal cells.

**Figure 5.**
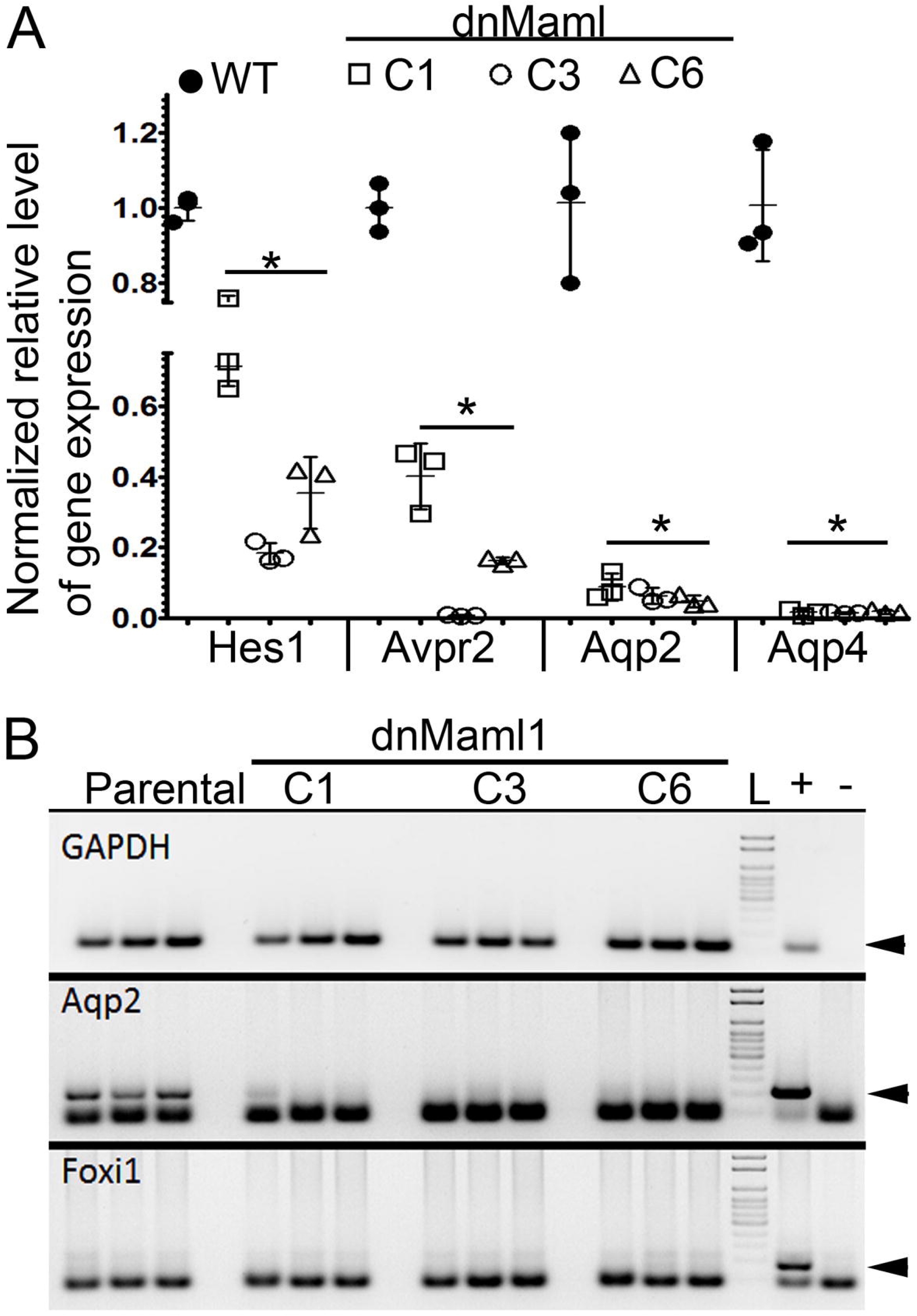
Suppression of Notch signaling in mature principal cells down regulates expression of principal cell specific genes without activating Foxi1 expression. **A.** Inhibition of Notch signaling by expressing dnMaml in mpkccdc14 principal cell line revealed reduced expression of Hes1 and reduced expression of principal cell genes as determined by RT-qPCR (* denotes p<0.05, two-tailed unpaired t-test; n=3 per cell line). **B.** Gel images of RT-PCR products reveals that Foxi1 expression is not detected in the parental or in the dnMaml expressing cells.

## Discussion

The patterning of the kidney collecting duct segments with principal cell types intermingled with intercalated cell types is hypothesized to involve Notch signaling mediated lateral inhibition (Fig.1). Here we have determined that among the family of Hairy/Enhancer of Split (HES) transcriptional repressors, which mediate Notch lateral inhibitory signal in other systems, Hes1 is critical for principal cell differentiation and maintenance of principal cell specific gene expression. Characterization of Hes1-mutant kidneys revealed a reduction in principal cell number and principal cell specific gene expression along with an increase in *Foxi1* expression and an increase in intercalated cells (Fig.2). This led us to test whether the main function of Hes1 is to repress *Foxi1* expression, by determining whether *Foxi1* inactivation rescues the kidney defects in mice with Hes1-deficient kidney collecting ducts. Interestingly, *Foxi1* inactivation rescues principal cell fate selection but does not restore the normal levels of principal cell specific gene expression (Fig. 3 and 4). This is evident with the increase in number of principal cells in *Cdh16->Cre;Hes1*^*f/f*^;Foxi1^−/−^ kidneys compared with *Cdh16->Cre;Hes1*^*f/f*^ kidneys (Fig.3), along with the continued occurrence of Aqp4-expressing cells without Aqp2 expression in the *Cdh16->Cre;Hes1*^*f/f*^; Foxi1^−/−^ kidneys (Fig.3). Additionally, the overall reduction in expression levels of *Aqp2* is restored back to normal levels in *Cdh16->Cre;Hes1*^*f/f*^; Foxi1^−/−^ kidneys but not the overall expression levels of *Aqp4* and *Elf5* (Fig.4). Even though not all principal cells in the *Cdh16->Cre;Hes1*^*f/f*^; Foxi1^−/−^ mouse kidneys express normal level of principal cell specific proteins, the increase in principal cell number is sufficient to rescue the urine concentrating defect and hydronephrosis observed in *Cdh16->Cre;Hes1*^*f/f*^ mice.

This is a significant finding as it suggests that by preventing *Foxi1* expression in mature kidney principal cells we should be able to prevent pathologies that are dependent on excessive conversion of mature principal cells into intercalated cells, such as during lithium treatment induced nephrogenic diabetes insipidus (NDI). Lithium, which is prescribed for management of bipolar disorders, does as a side effect cause kidney dysfunction that manifests initially as a urine concentrating defect. At the cellular level lithium triggers an increase in intercalated cell numbers and a reduction in principal cell numbers. We have recently determined by lineage tracing of principal cells in mature mouse kidney collecting ducts that lithium triggers the conversion of labeled principal cells into intercalated cells [34]. It will be important to determine if *Foxi1* inactivation can prevent lithium induced urine concentrating defects.

In conclusion, among the Notch signaling downstream targets, Hes1 is a critical component that ensures collecting duct cell type patterning by primarily repressing *Foxi1*. There is likely to be additional roles for Hes1 apart from repressing *Foxi1*, since *Foxi1* inactivation cannot rescue the defect in maintenance of expression of all principal cell genes in the Hes1-deficient kidney collecting ducts. It is likely that additional factors are repressed by Hes1 to ensure maintenance of the mature principal cell state and function. Alternatively, there are reports that Hes1 can function as a transcriptional activator in other biological contexts [35, 36], and hence Hes1 may act as a transcriptional activator in mature principal cells by directly activating principal cell gene transcription.

## Acknowledgements

Research reported here was supported by a grant from NIDDK of NIH under award number R01DK106135. We thank the Histology and Imaging Core, and Functional Genomics and Bioinformatics cores at Sanford Research for technical assistance. These Cores at Sanford Research were supported by NIH grants P20GM10358, P20GM121341 and P20GM103620.

## Author Contributions

M. Mukherjee, M. Janga, E. Fogarty, and J. DeRiso performed experiments, and analyzed the results; M. Mukherjee and J. DeRiso wrote the methods sections; K. Surendran designed experiments, helped analyze the results and wrote the manuscript; and all authors contributed to manuscript editing and revisions.

## Competing Interest

The authors declare no competing interests.

## Supplementary Figures

**Figure S1. Foxi1-/- mouse kidneys do not express intercalated cell specific markers expect for carbonic anhydrase II (CAII).** Analysis of post-natal day 0 kidneys from wild type (WT) mice (A&C) versus Foxi1-/- mice (B&D). The intercalated cell markers Atp6v1b1 and Ae1 are expressed in renal epithelial cells of cortically located tubules (arrows in A) but not in cortically located tubules of Foxi1-/- mouse kidneys (B). Smaller interstitial cells (asterisks in A &B) are labeled with the Ae1 antibody in both WT and Foxi1-/- kidneys. CAII expressing renal epithelial cells are present in WT kidneys of P0 (arrows in C) and 7 week (arrows in E) old mice that are located next to Aqp2+ cells within medullary collecting duct segments. CAII expression is not detectable in medullary collecting duct segments in P0 Foxi1-/- kidneys (D) but is evident in Aqp2+ cells of medullary collecting duct segments of 7 week old Foxi1-/- kidneys (F). All scale bars represent 10μm.

**Figure S2. *Foxi1* inactivation rescues principal cell fate selection deficiency but not the reduced principal cell gene expression in *Cdh16-Cre;Hes1f/f* mouse kidneys.** Adult mouse kidneys from Hes1f/f controls (A & D) contain c-Kit expressing intercalated cells (asterisks in A&D)) intermingled among Aqp3+ (A) and Aqp2+;Aqp4+ (D) principal cells. Inactivation of Hes1 in Cdh16-Cre;Hes1f/f adult mouse kidneys (B&E) results in increased number of c-Kit expressing intercalated cells (asterisks in B&E) and reduced expression of Aqp3 (arrow in B) or Aqp2 and Aqp4 (arrow in E) in some cells that selected the principal cell fate. Combined inactivation of Foxi1 and Hes1 in *Cdh16-Cre;Hes1f/f;Foxi1-/-* mouse kidneys (C &E) result in no c-Kit expressing renal epithelial cells intermingled among Aqp3+ cells (C). Some cells lack both c-Kit and Aqp3 expression (arrows in C) and likely are principal cells with reduced Aqp3 expression. Collecting duct cells with reduced level of basolateral Aqp4 and/or reduced apical Aqp2 expression (arrows in F & F’) are common in *Cdh16-Cre;Hes1f/f;Foxi1-/-* mouse kidneys. All scale bars represent 10μm.

## References

1. Costantini, F. and R. Kopan, Patterning a complex organ: branching morphogenesis and nephron segmentation in kidney development. Dev Cell, 2010. 18(5): p. 698–712.

2. Brown, D., P. Weyer, and L. Orci, Vasopressin stimulates endocytosis in kidney collecting duct principal cells. Eur J Cell Biol, 1988. 46(2):p. 336–41.

3. Brown, D. and T. Kumpulainen, Immunocytochemical localization of carbonic anhydrase on ultrathin frozen sections with protein A-gold. Histochemistry, 1985. 83(2): p. 153–8.

4. Bastani, B., Immunocytochemical localization of the vacuolar H(+)-ATPase pump in the kidney. Histol Histopathol, 1997. 12(3): p. 769–79.

5. Kim, J., C.C. Tisher, and K.M. Madsen, Differentiation of intercalated cells in developing rat kidney: an immunohistochemical study. Am J Physiol, 1994. 266(6 Pt 2): p. F977–90.

6. Emmons, C. and I. Kurtz, Functional characterization of three intercalated cell subtypes in the rabbit outer cortical collecting duct. J Clin Invest, 1994. 93(1): p. 417–23.

7. Fejes-Toth, G., et al., Differential expression of AE1 in renal HCO3-secreting and -reabsorbing intercalated cells. J Biol Chem, 1994. 269(43): p. 26717–21.

8. Brown, D. and S. Breton, Mitochondria-rich, proton-secreting epithelial cells. J Exp Biol, 1996. 199(Pt 11): p. 2345–58.

9. Brown, D., V. Ilic, and L. Orci, Rod-shaped particles in the plasma membrane of the mitochondria-rich cell of amphibian epidermis. Anat Rec, 1978. 192(2): p. 269–75.

10. Janicke, M., T.J. Carney, and M. Hammerschmidt, Foxi3 transcription factors and Notch signaling control the formation of skin ionocytes from epidermal precursors of the zebrafish embryo. Dev Biol, 2007. 307(2): p. 258–71.

11. Grimm, P.R., et al., Integrated compensatory network is activated in the absence of NCC phosphorylation. J Clin Invest, 2015. 125(5): p. 2136–50.

12. Quigley, I.K., J.L. Stubbs, and C. Kintner, Specification of ion transport cells in the Xenopus larval skin. Development, 2011. 138(4): p. 705–14.

13. Blomqvist, S.R., et al., Distal renal tubular acidosis in mice that lack the forkhead transcription factor Foxi1. J Clin Invest, 2004. 113(11): p. 1560–70.

14. Jeong, H.W., et al., Inactivation of Notch signaling in the renal collecting duct causes nephrogenic diabetes insipidus in mice. J Clin Invest, 2009. 119(11): p. 3290–300.

15. Guo, Q., et al., Adam10 Mediates the Choice between Principal Cells and Intercalated Cells in the Kidney. J Am Soc Nephrol, 2015. 26(1): p. 149–59.

16. Kopan, R. and M.X. Ilagan, The canonical Notch signaling pathway: unfolding the activation mechanism. Cell, 2009. 137(2): p. 216–33.

17. De Strooper, B., et al., A presenilin-1-dependent gamma-secretase-like protease mediates release of Notch intracellular domain. Nature, 1999. 398(6727): p. 518–22.

18. Steiner, H., et al., A loss of function mutation of presenilin-2 interferes with amyloid beta-peptide production and notch signaling. J Biol Chem, 1999. 274(40): p. 28669–73.

19. Kitagawa, M., et al., A human protein with sequence similarity to Drosophila mastermind coordinates the nuclear form of notch and a CSL protein to build a transcriptional activator complex on target promoters. Mol Cell Biol, 2001. 21(13): p. 4337–46.

20. Grassmeyer, J., et al., Elf5 is a principal cell lineage specific transcription factor in the kidney that contributes to Aqp2 and Avpr2 gene expression. Dev Biol, 2017. 424(1): p. 77–89.

21. Vidarsson, H., et al., The forkhead transcription factor Foxi1 is a master regulator of vacuolar H-ATPase proton pump subunits in the inner ear, kidney and epididymis. PLoS One, 2009. 4(2): p. e4471.

22. Kurth, I., et al., The forkhead transcription factor Foxi1 directly activates the AE4 promoter. Biochem J, 2006. 393(Pt 1): p. 277–83.

23. Yang, T., et al., Transcriptional control of SLC26A4 is involved in Pendred syndrome and nonsyndromic enlargement of vestibular aqueduct (DFNB4). Am J Hum Genet, 2007. 80(6): p. 1055–63.

24. Kageyama, R., et al., Dynamic Notch signaling in neural progenitor cells and a revised view of lateral inhibition. Nat Neurosci, 2008. 11(11): p. 1247–51.

25. Heitzler, P. and P. Simpson, The choice of cell fate in the epidermis of Drosophila. Cell, 1991. 64(6): p. 1083–92.

26. Bray, S., Notch signalling in Drosophila: three ways to use a pathway. Semin Cell Dev Biol, 1998. 9(6): p. 591–7.

27. Stratman, J.L., W.M. Barnes, and T.C. Simon, Universal PCR genotyping assay that achieves single copy sensitivity with any primer pair. Transgenic Res, 2003. 12(4): p. 521–2.

28. Bens, M., et al., Corticosteroid-dependent sodium transport in a novel immortalized mouse collecting duct principal cell line. J Am Soc Nephrol, 1999. 10(5): p. 923–34.

29. Hrabe de Angelis, M., J. McIntyre, 2nd, and A. Gossler, Maintenance of somite borders in mice requires the Delta homologue DII1. Nature, 1997. 386(6626): p. 717–21.

30. Piscione, T.D., M.Y. Wu, and S.E. Quaggin, Expression of Hairy/Enhancer of Split genes, Hes1 and Hes5, during murine nephron morphogenesis. Gene Expr Patterns, 2004. 4(6): p. 707–11.

31. Chen, L. and Q. Al-Awqati, Segmental expression of Notch and Hairy genes in nephrogenesis. Am J Physiol Renal Physiol, 2005. 288(5): p. F939–52.

32. Imayoshi, I., et al., Hes genes and neurogenin regulate non-neural versus neural fate specification in the dorsal telencephalic midline. Development, 2008. 135(15): p. 2531–41.

33. Shao, X., S. Somlo, and P. Igarashi, Epithelial-specific Cre/lox recombination in the developing kidney and genitourinary tract. J Am Soc Nephrol, 2002. 13(7): p. 1837–46.

34. Mukherjee, M., et al., Endogenous Notch Signaling in Adult Kidneys Maintains Segment-Specific Epithelial Cell Types of the Distal Tubules and Collecting Ducts to Ensure Water Homeostasis. J Am Soc Nephrol, 2019. 30(1): p. 110–126.

35. Sugita, S., et al., Transcription factor Hes1 modulates osteoarthritis development in cooperation with calcium/calmodulin-dependent protein kinase 2. Proc Natl Acad Sci U S A, 2015. 112(10): p. 3080–5.

36. Ju, B.G., et al., Activating the PARP-1 sensor component of the groucho/TLE1 corepressor complex mediates a CaMKinase IIdelta-dependent neurogenic gene activation pathway. Cell, 2004. 119(6): p. 815–29.

