## Supplementary figures and images for "*Foxi1* inactivation rescues loss of principal cell fate selection in *Hes1*-deficient kidneys but does not ensure maintenance of principal cell gene expression"

### Figure S1

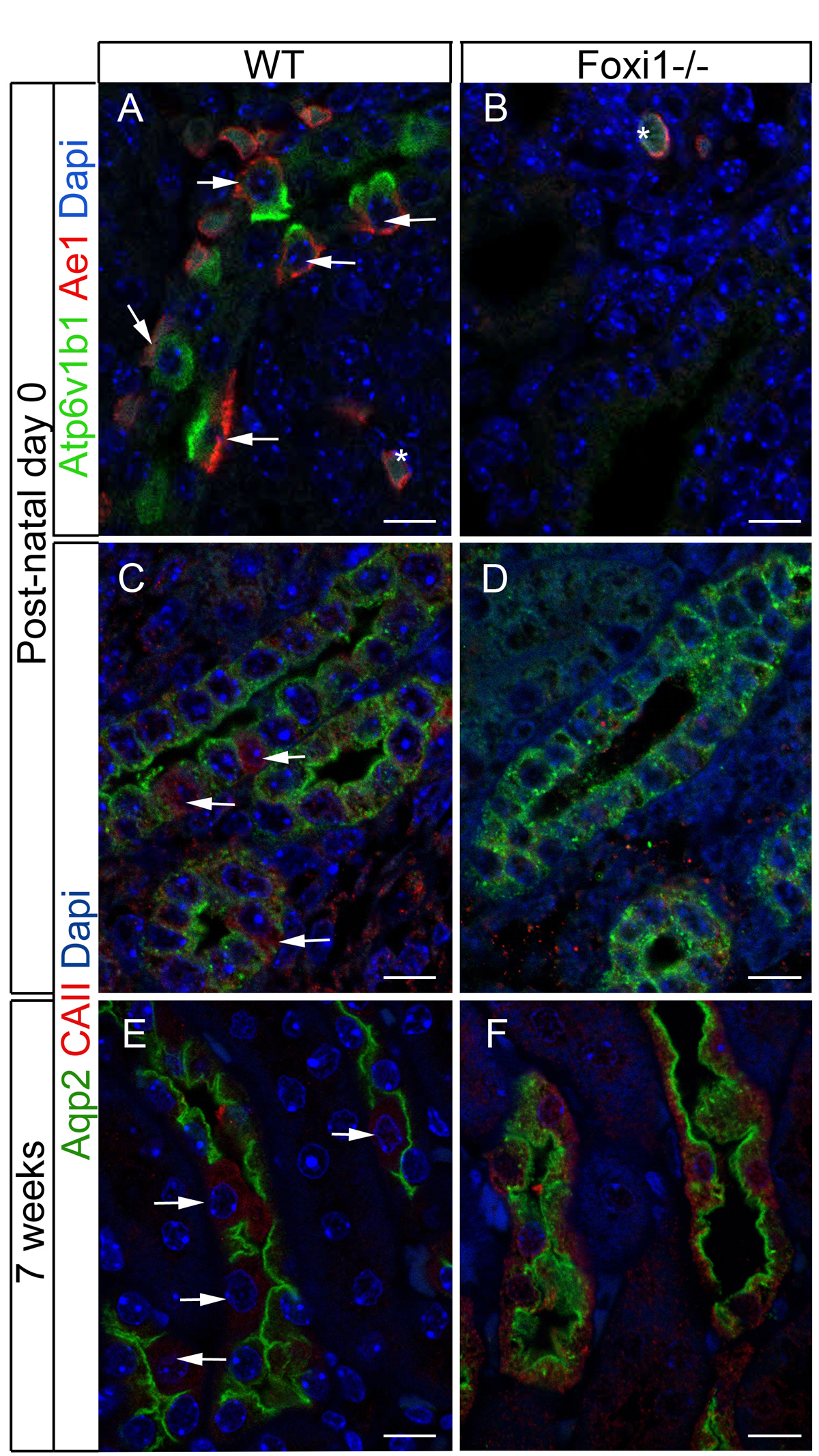

### Figure S2

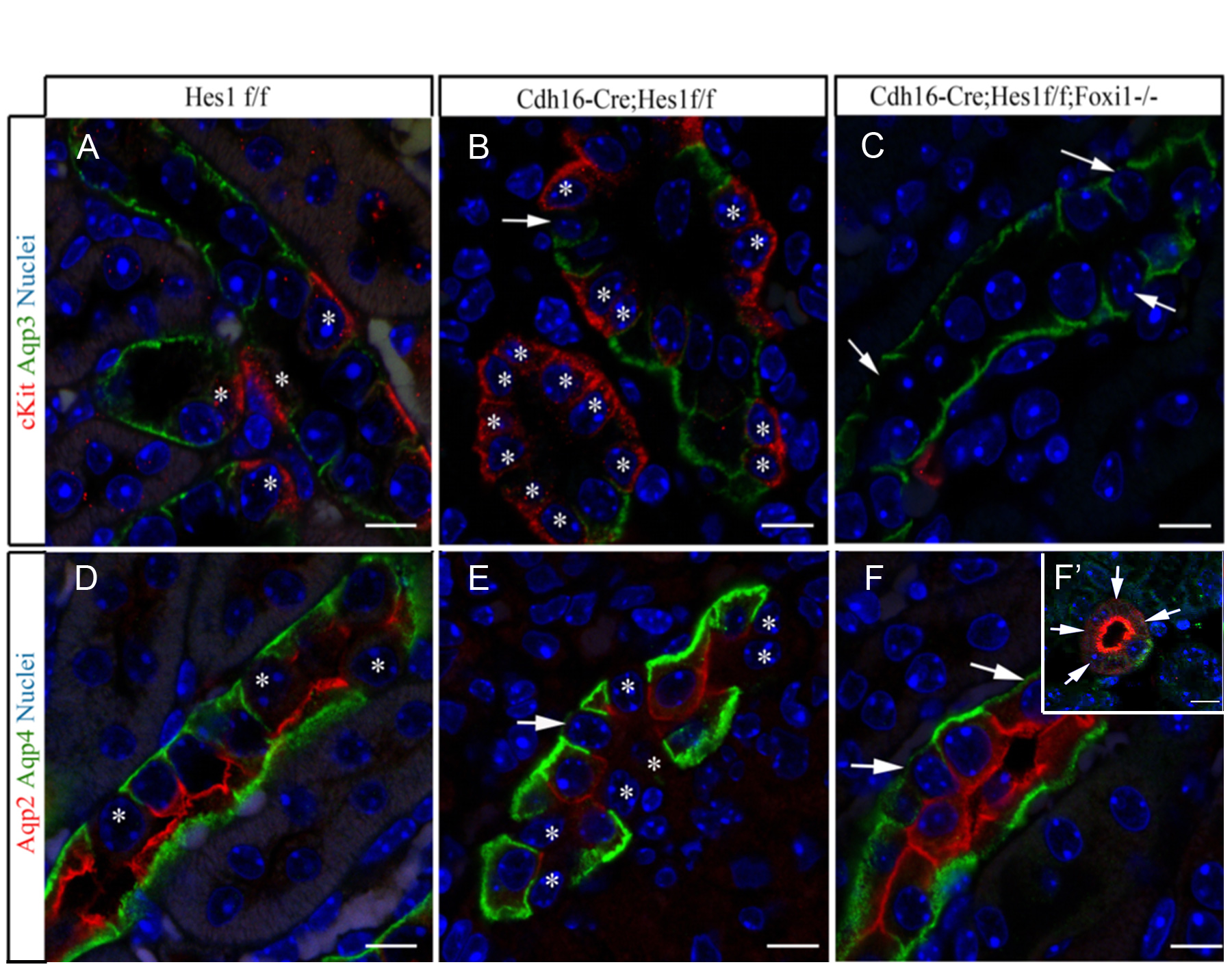
